# Testing two competing hypotheses for Eurasian jays’ caching for the future: planning versus compensatory caching

**DOI:** 10.1101/2020.07.13.200063

**Authors:** Piero Amodio, Johanni Brea, Benjamin G. Farrar, Ljerka Ostojic, Nicola S. Clayton

## Abstract

Previous research reported that corvids preferentially cache food in a location where no food will be available or cache more of a specific food in a location where this food will not be available. Here, we consider possible explanations for these prospective caching behaviours and directly compare two competing hypotheses. The Compensatory Caching Hypothesis suggests that birds learn to cache more of a particular food in places where that food was less frequently available in the past. In contrast, the Future Planning Hypothesis suggests that birds recall what-when-where features of specific past events to predict the future availability of food. We designed a protocol in which the two hypotheses predict different caching patterns across different caching locations such that the two explanations can be disambiguated. We formalised the hypotheses in a Bayesian model comparison and tested this protocol in two experiments with one of the previously tested species, namely Eurasian jays. Consistently across the two experiments, the observed caching pattern did not support either hypothesis; rather it was best explained by a uniform distribution of caches over the different caching locations. Future research is needed to gain more insight into the cognitive mechanism underpinning corvids’ caching for the future.

## 1 Introduction

Over the last two decades, an increasing number of studies in non-human animals have challenged the view that future planning abilities are uniquely human (primates: [1, 2, 3, 4, 5]; corvids: [6, 7, 8, 9]; rodents: [10, 11], although see [2]). Within corvids, California scrub-jays (*Aphelocoma californica*) and Eurasian jays (*Garrulus glandarius*) have been shown to adapt their caching behaviour according to future needs. Experimental studies reported that these jays cached more food in locations where food was going to be absent in the following trial, or preferentially cached a specific type of food in locations where that food was not going to be available in the following trial ([6, 7, 9, 12]).

The jays’ performance in these studies is in line with the Future Planning Hypothesis: at the time of caching, jays may have recalled the ‘what-when-where’ features of past events – the availability of a specific food in a given location at a specific time – and cached food that will maximise their future outcome, namely, at the time when they would retrieve their caches. It has been suggested that jays may be able to do so even when their motivational state at the time of caching differs from their motivational state at retrieval, such that their caching decision is not based on their current desire for the food in question [6, 7]. One manner in which an individual’s current motivation can be manipulated is through sating an individual on a specific food. This procedure subsequently reduces the individual’s motivation for eating that specific food (but not different kinds of food), a phenomenon known as ‘specific satiety’[13, 14]. Utilising this phenomenon in their experimental manipulations, Cheke and Clayton [6] and Correia et al. [7] investigated the jays’ caching behaviour when their motivational state at the time of caching and at the time of retrieval differed, by using a within- and a between-subjects design, respectively. The authors report that jays cached preferentially a type of food that was not going to be available in the future even when they had a low desire toward that food at the time of caching. Additional support for the Future Planning Hypothesis comes from experiments reporting that jays may possess ‘episodic-like memory’ [15], i.e., that they can represent and subsequently recall information about what happened in a specific event, where it took place, and when it occurred in an integrated manner [16]. These findings provide support for the Future Planning Hypothesis because i) this hypothesis hinges on the ability to recall ‘what-where-when’ features of specific past events [17], ii) episodic-like memory and future planning are thought to be based on the same cognitive machinery [18, 19] and, iii) show the same developmental trajectory in young children [20].

However, it has been argued that the jays’ performance in these caching experiments cannot be taken as evidence of future planning abilities because the experimental designs do not exclude alternative interpretations. The two predominant alternative explanations that have been raised are the Mnemonic Association Hypothesis and the Compensatory Caching Hypothesis. Importantly, these two hypotheses are not necessarily mutually exclusive; rather, they may describe two parallel cognitive processes. Indeed, neither of these hypotheses can provide an alternative interpretation for all published caching experiments, but only to a specific subset of the studies. However, together they can explain all of the currently reported results.

The Mnemonic Association Hypothesis ([17]; see also [6]) suggests that jays’ strategic caching behaviour may be the result of long-delay learning, guided by what the birds have learned at the time of cache recovery [21]. At the time of the outcome (e.g. at retrieval of previously made caches), jays may have recalled their previous actions, and thus created a positive association with the specific action that had resulted in a more beneficial outcome, i.e. the action of caching a specific type of food in that specific location at a specific time. This mechanism hinges on episodic-like memory because the retrieval of the ‘what-when-where’ features of this specific past episode is essential to develop such a positive association between the outcome experienced in the present and the action that was performed in the past and that led to that outcome. Accordingly, in the studies conducted by Correia et al. [7], de Kort et al. [12], and Cheke and Clayton [6], jays may have – during the second and third caching session – reiterated the actions which had led to the more beneficial outcome in the first recovery session, thereby solving the tasks without pre-experiencing future scenarios.In a similar vein, this mechanism may also account for the results of Clayton et al. [22]’s study, in which it was reported that scrub-jays could learn to refrain from caching a specific type of food when, consistently across multiple trials, that specific food was always degraded at the time of recovery, due to experimental manipulation. Thus, although both the Mnemonic Association Hypothesis and the Future Planning Hypothesis postulate ‘what-where-when’ memory, the former explains the jays’ strategic caching as the result of a learning process at the time of cache retrieval, while the latter explains it as the result of a prospective process at the time of caching.

Crucially however, while the Future Planning Hypothesis can explain the data from all experiments that have been published, the Mnemonic Association Hypothesis cannot provide an explanation for the results reported by Raby et al.[9]. Here, on six consecutive days, California scrub-jays had access to either compartment A or B during testing, with the availability of food during the first two hours being dependent on the compartment the bird was in (Figure 1). Food was freely available for the rest of the day. In the evening of the sixth day the jays were allowed to cache for the first time: the authors reported that jays cached more food in the compartment where no food was available on the previous mornings than in the compartment where it was available [9]. In a follow-up experiment, each of the two testing compartments was associated with a specific type of food (e.g. food X available in compartment A, food Y available in compartment B; Figure 1). The authors reported that, on the test trial, the jays preferred to cache the food that had not been available in that compartment as opposed to the food that had been previously available there [9]. These results appear to be consistent with the Future Planning Hypothesis: jays may have adjusted their caching strategy according to their expectation of the specific future event, thereby ensuring that on the hypothetical following day, food will be available in the compartment where they experienced hunger on the previous days (in Experiment 1) or that both types of foods will be present in the two compartments (in Experiment 2). By contrast, these results cannot be explained by the Mnemonic Association Hypothesis. Jays were given the opportunity to cache only once, namely at the test, such that their caching pattern could not have resulted from long-delay learning. In particular, jays could not have adjusted their caching behaviour according to the outcome of a previous caching event experienced at retrieval, because no such previous caching event and related retrieval event existed [17]. Nevertheless, it has been argued that the jays’ performance in this study may not entail any kind of prospection ability. Instead, the jays may have had ‘a propensity to cache a particular food type in a given location that differs from the foods that have been previously associated with that location, a strategy that would provide more uniform distribution of resources’ (Shettleworth personal communication, as cited in [17], p. 90; see also [23, 24]). Support for this idea is, to some extent, provided by field observations: wild American crows (*Corvus brachyrhynchos*) foraging at walnut trees were never observed caching nuts in proximity of the fruiting trees but carried the items up to 2km away before caching them [25]. This argument, henceforth referred to as the Compensatory Caching Hypothesis, appears to bear high ecological relevance also in the case in which two different types of food are available. As pointed out by Shettleworth [26], ‘For an animal that caches different types of items (and […] can remember what it cached where), a strategy of distributing items of each type as widely as possible would help to defeat predators that might raid just one of those types’ (p. 393). Since caching behaviour is considered to have played a key role in the evolution of corvid cognition [27, 28], it appears plausible that, to protect their caches, these birds may possess predispositions such as those assumed by the Compensatory Caching Hypothesis.

**Figure 1:**
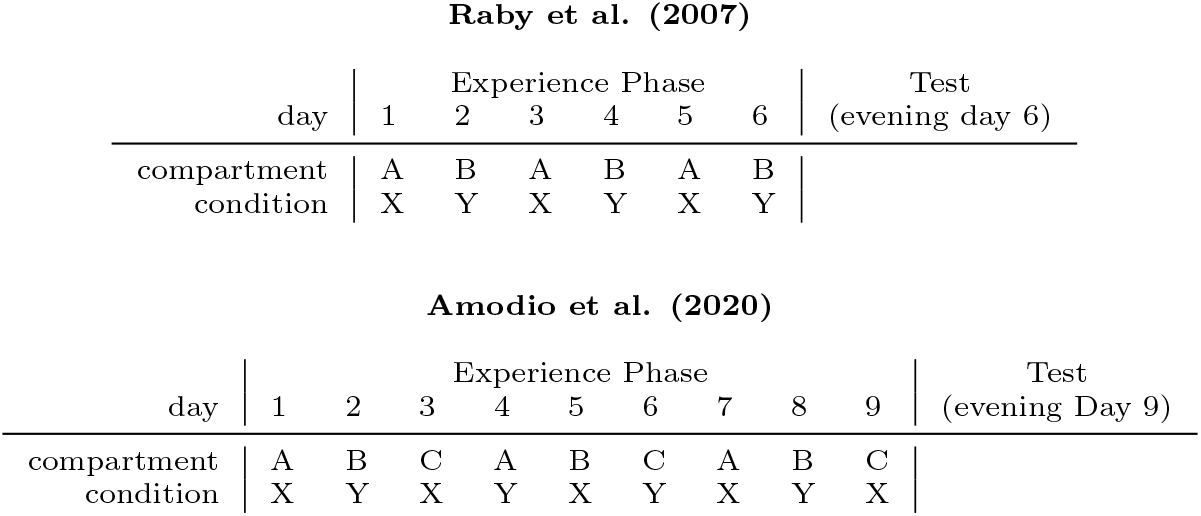
Experimental protocols. On training days, the birds could access one of the compartments (A, B or C) under condition X or Y, where X could mean that no food is provided or only food of a certain type, and Y could mean that food is provided or only food of another type. In the protocol of Raby et al. (2007), each compartment is always experienced under the same condition. In the experiment reported here, each compartment is experienced under both conditions.

To reach firmer conclusions on whether caching behaviour in corvids entails prospection abilities, it is essential to test the Future Planning Hypothesis against each of the two alternative interpretations that are currently available. As the Mnemonic Association Hypothesis and the Compensatory Caching Hypothesis may describe two complementary processes, these hypotheses can be investigated independently. Here we present a first step in this direction, by focusing on the Complementary Caching Hypothesis. We propose a mechanism that may underpin jays’ propensity to distribute caches such that across different locations resources are uniformly distributed and empirically test the Complementary Caching Hypothesis against the Mental Time Travel Hypothesis in Eurasian jays.

We suggest that jays may evaluate the suitability of a given location as a caching site for a specific type of food by attributing to it location-specific ‘weights’ according to the perceived availability of that food in that location. For instance, if a jay is given the possibility to access three potential caching locations (A, B, C), in which no food was experienced, then it may attribute equal weights to each location (wA, wB, wC). Therefore, when allowed to cache, the jay may distribute caches in comparable quantities across the locations. However, if the jay subsequently experiences a given food as being available in a given location, for example at location A, then the location-specific weights may be updated accordingly, i.e. wA would be reduced in comparison to wB and wC. As a result, when allowed to cache, the jay may concentrate its caches in the two locations associated with higher weights for this food (e.g. locations B and C), thus achieving a more uniform distribution of resources. This idea is similar to that proposed by Hampton and Sherry [29] as a possible mechanism employed by black-capped chickadees for re-using caching locations according to the probability of recovering previously hidden items.

Building on Raby et al. [9]’s study, we devised a paradigm to differentiate between the Compensatory Caching and the Future Planning Hypotheses (Figure 1). Eurasian jays first received an experience phase over nine consecutive days. On each day, jays had access to one of three compartments. In addition, we manipulated whether and what food jays received in each compartment on each day. Critically, location and food availability were changed on a 3-day cycle and on a 2-day cycle, respectively (Figure 1). Thus, the three compartments were differently associated with (different) food, i.e., each compartment was associated either once or twice with each type of food. Subsequently, a single test trial was conducted in which jays could freely cache both foods in all three locations at the same time. This design ensured that the two competing hypotheses predict opposite caching patterns. In Experiment 1, the location was changed on a 3-day basis and which of two foods (food X or food Y) was received was changed on a 2-day basis. In Experiment 2, we decreased the cognitive load of the experiment, such that only one food type was available. Consequently, the location was changed on a 3-day basis and whether or not food was available was changed on a 2-day basis.

## 2 General Methods

### 2.2 Subjects

Nine Eurasian jays of both sexes (all born in 2007) participated in the study: Caracas (m), Lima (m), Lisbon (m), Dublin (m), Rome (f), Jerusalem (f), Wellington (f), Washington (f) and Quito (f). All birds were tested in both Experiment 1 and Experiment 2, except one individual (Jerusalem) that was tested only in Experiment 1. This bird was euthanized due to sickness unrelated to the behavioural tests prior to the start of Experiment 2. Birds were housed as a group in a large outdoor aviary (20 × 10 × 3 m) at the Sub-Department of Animal Behaviour, University of Cambridge, Madingley. The sample size included all birds of the colony of jays available to us for this study. Two other colonies that are housed at the same site were not tested because these birds were being used in other studies at that time; thereby it was not possible to install the set-up required for the experiments reported here in their indoor testing compartments. Outside of testing the birds had *ad libitum* access to their maintenance diet, which consisted of vegetables, eggs, seed and fruits. Water was available at all times. All procedures were approved by the University of Cambridge Animal Ethics Review Committee.

### 2.2 Experimental Set-up

The experimental set-up comprised three indoor test compartments (labelled A, B, C) that were accessible from an equidistant middle compartment (Figure 2). All four compartments (1 × 1 × 2 m) were delimited by mesh walls and contained a suspended wooden platform (1 × 1 m). Birds could access the set-up from a trap-window connecting the outside aviary with the middle compartment, and from there they could reach each test compartment through an opening in the mesh wall. Designated areas in the mesh had transparent Perspex windows in them and openings were available to the birds when the windows were raised. Whilst standing outside the compartment, an experimenter could raise or lower the windows using ropes attached to the windows.

**Figure 2:**
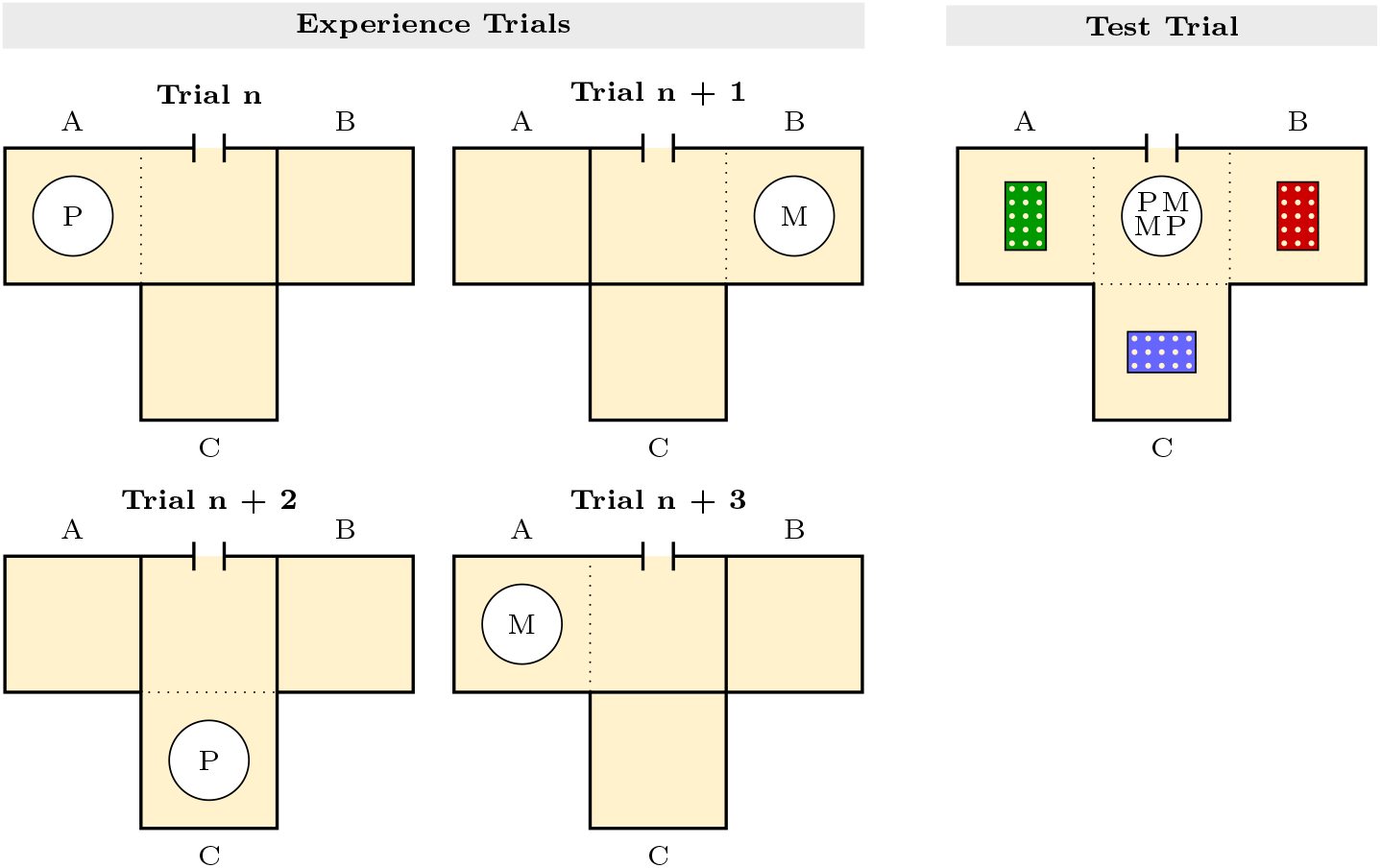
Top view of the experimental set-up of Experiment 1. Birds could access the three test compartments (A, B, C) from a middle compartment. On each experience trial (Left), the bird had access only to one test compartment (e.g. A in Trial n; B in Trial n+1), and one type of food in powdered form (e.g. Peanuts (P) in Trial n; Macadamia nuts (M) in Trial n+1). The compartment that was accessible and food available rotated respectively on a 3-days cycle and on a 2-days cycle. On the test trial (Right), a bowl containing items of both types of food was placed in the middle compartment and caching trays were placed in all test compartment. Birds could freely move and cache in all compartments.

## 3 Experiment 1

In this experiment, jays had access to one of three compartments in which a specific food (food X or food Y) was available for nine days. The location and the type of food available on each day changed on a 3-day cycle and on a 2-day cycle, respectively (Figure 1). Consequently, the jays experienced the three compartments as being differentially associated with each specific food. The experiment was conducted from October to December 2017. LO, BF and PA were the experimenters.

### 3.1 Pre-Test

The test procedure required that jays in principle eat powdered food in sufficient quantity to develop specific satiety. Note that it was essential to use food in powdered form so that jays could not cache inside the experimental set-up during the experience phase. To ensure that jays would meet these requirements, a pre-test was conducted. Each bird received two pre-test trials in total. The birds’ maintenance diet was removed from the aviary approximately 1.5 h prior to the start of each trial. A bird was tested in visual isolation from the rest of the group, namely in an indoor compartment that was not part of the experimental set-up. Trials were conducted on different days and involved a pre-feeding phase followed by a test phase. On each trial, the bird was first provided with a bowl containing 50.0 g of powdered food (either peanuts or macadamia nuts) and given the opportunity to eat for 15 minutes. Following this pre-feeding phase, the powdered food bowl was removed and the bird received two bowls, one containing 50 whole peanuts and the other containing 50 macadamia nut halves. The bird could freely eat and manipulate (and thus also cache in the compartment) both types of food during the test phase for 15 minutes. At the end of each trial, we recorded: i) the amount of powdered food taken out of the bowl during the pre-feeding phase, and; ii) the number of items of both foods that were taken from the bowls in the testing phase. Each bird experienced both types of food in powdered form in the pre-feeding phase, one in each trial. The order in which the two types of foods were provided to the birds during the pre-feeding phase was counterbalanced across birds. To pass the pre-test, a bird was required to i) take at least one peanut and one macadamia nut from the respective bowls across the two trials, and ii) exhibit an eating pattern that was numerically in line with specific satiety across the two trials, i.e. to show a relative preference for the non-pre-fed food when both trials were considered. For instance, a smaller number of peanuts after being pre-fed peanuts than after being pre-fed macadamia nuts would be in line with specific satiety. If a bird did not meet the criteria on the first pair of trials, it was re-tested a second time.

### 3.2 Familiarisation

After the pre-test, birds were familiarised with the experimental set-up in one trial. A bird was individually given access to the middle compartment and allowed to explore all three testing compartments at the same time. At this stage, the windows to all compartments were kept open. Three items of a high value food that was not used as a test food in the later test (i.e., wax moth larvae, *Galleria mellonella*) were placed in each test compartment to entice the bird to explore the different areas. The bird inside the set-up was monitored remotely through a CCTV camera system. The duration of the familiarisation was not the same for all birds: a bird was released back into the aviary through the trap-window in the middle compartment after it has eaten in all compartments. However, if a bird showed no motivation for the food or otherwise looked as if they were not comfortable in the set-up (e.g., flying across compartments for a prolonged time), the trial was ended due to welfare reasons and the bird released into the outdoor aviary. Thus, no standardised duration was set for the trials. The decision regarding how long a bird was inside the set-up was made by the experimenter on a case-by-case basis. To successfully complete the familiarisation, birds were required to eat at least one food item in each test compartment. In case a bird did not reach this criterion, the familiarisation trial was repeated up to two times (i.e., birds could receive a maximum of three familiarisation trials).

### 3.3 Test

The experiment consisted of an experience phase involving nine experience trials, followed by one test trial. In the experience phase birds received one trial per day on nine consecutive days. Each trial was conducted in the morning, after the birds’ maintenance diet had been removed from the aviary for approximately 1.5 h to ensure that birds were mildly hungry and thus motivated for food. On each trial birds were individually allowed to access the experimental set-up through the trap-window in the middle compartment and given access only to one of the three test compartments. A bowl containing powdered food (50.0 g of peanuts or 50.0 g of macadamia nuts) was placed on the suspended platform in the accessible test compartment. Food in powdered form was used to prevent the bird from caching in the test compartments during the experience phase. The bird could freely eat and move between the middle compartment and the accessible test compartment for 15 minutes. At the end of a trial the bird was released into the outside aviary through the trap-window in the middle compartment. The order in which the different birds were tested was kept the same across all trials (i.e. the same bird was the first bird to be the tested on all days, and so on). The location and the type of food experienced by the birds on each trial changed according to two different patterns (Figure 1 and Figure 2). The test compartment that was accessible rotated on a 3-day basis, such that, for example, the birds that had access to compartment A on trial 1 experienced the test compartments in the following order across the nine trials: A-B-C-A-B-C-A-B-C. The type of food available rotated on a 2-day basis, i.e. the same food was provided every other trial. As a result, all test compartments were experienced by each bird three times, and differed in the frequency in which they were associated with a specific type of food. In particular, two compartments were associated twice with one type of food and once with the alternative food, whereas the third compartment was characterised by the opposite pattern. The compartment that was accessible and the type of food available on trial 1 were counterbalanced across birds.

The test phase was conducted on the day of the last experience trial (9th day), approximately 3h after the experience phase. Again, birds were individually given access to the experimental set-up. All test compartments were accessible at this stage, and each contained one caching tray (5 × 3 pots filled with sand). A bowl containing 25 peanuts and 25 macadamia nut halves was placed in the middle compartment. Inside the bowl, food items were arranged so that both types of food were equidistantly positioned from each opening between the middle compartment and a test compartment (Figure 2). Birds could eat and cache both types of food in all test compartments for 15 minutes. At the end of the trial birds were released to the outdoor aviary through the middle compartment. Birds were not given the opportunity to retrieve their caches.

### 3.4 Predictions

According to the Compensatory Caching Hypothesis, jays should cache a similar overall number of items in each of the three test compartments but with different proportions of the particular types of food. Specifically, they should exhibit a preference for caching in each compartment the food that was – during the experience phase – less frequently experienced in that specific compartment (Figure 3 bottom left). In contrast, the Future Planning Hypothesis predicts that birds should exhibit a preference for caching according to which location they would have access on the future days (i.e. hypothetical additional trials equivalent to past experience trials) and according to what food would be available there. Two possible, complementary, patterns would support the Future Planning Hypothesis: jays could either provision only for the nearest future event (hypothetical trial 10) or for all three trials that they could provision for at that time (hypothetical trials 10, 11, and 12). According to Prediction 1, jays should cache more items of the food that they expect would not be available and only in the compartment that they expect would be accessible on the hypothetical trial 10 (Figure 3 top left). By contrast, according to Prediction 2, jays should distribute their caches across all three compartments but cache more of a particular food in the specific compartment where they expect that it would not to be provided the next time that they have access to it (hypothetical trials 10, 11, and 12; Figure 3 centre left). Importantly, these two predictions of the Future Planning Hypothesis differ only in whether caching is expected in the two compartments that correspond to the hypothetical days 11 and 12.

**Figure 3:**
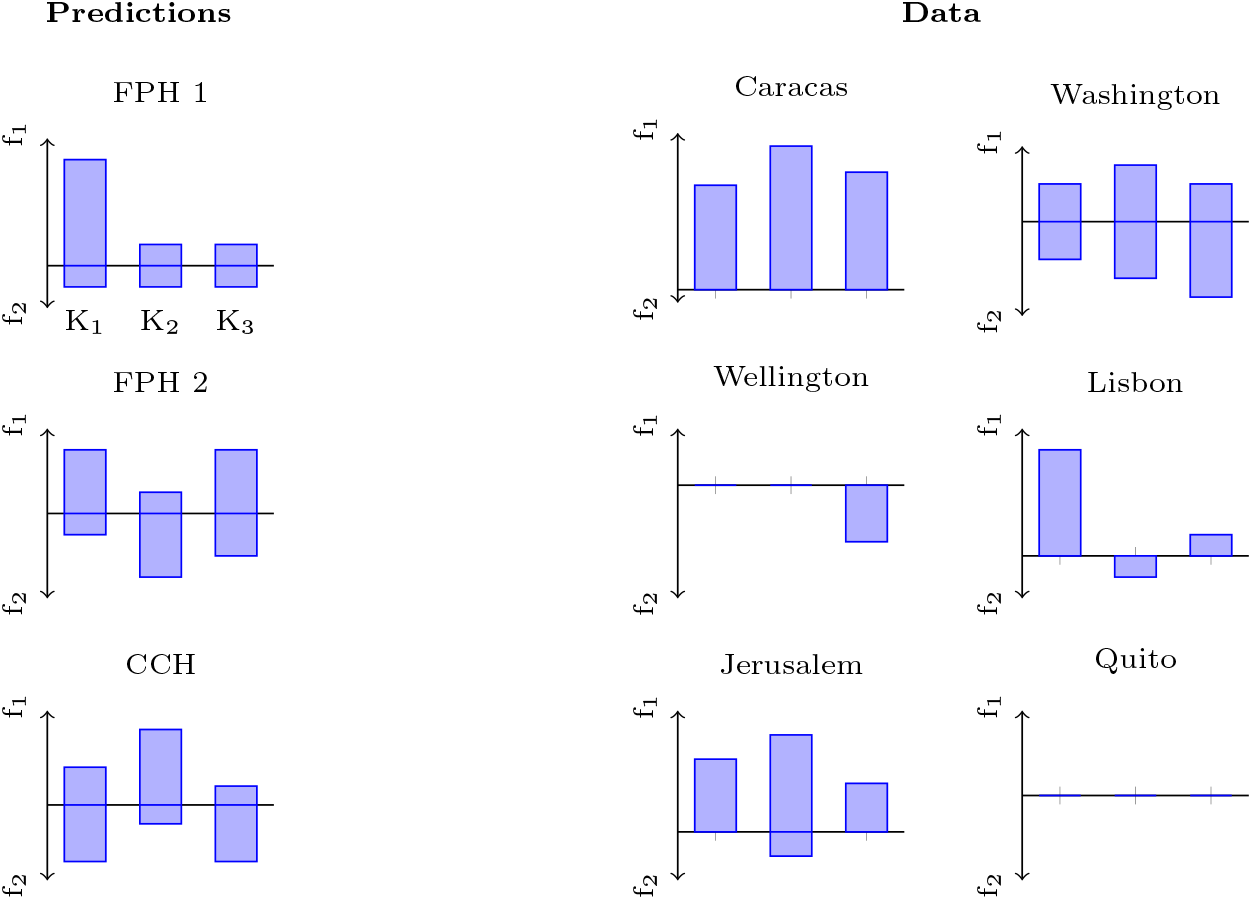
Individual caching patterns allow no conclusion. If a bird caches according to the Future Planning Hypothesis (FPH 1), it will provision only for the next day, where it expects to be in compartment K1 with food of type f2 available; thus it would cache food f1 in K1. If it also takes into account subsequent days (FPH 2), it will distribute the caches to complement the food type it expects to find in the respective compartment. If the bird caches according to the Compensatory Caching Hypothesis (CCH), the bird should cache less of food type f in compartments K, where it has previously encountered many items of food type f. The actual numbers in the predictions serve illustration purposes only. The hypotheses allow only qualitative predictions, e.g. the Compensatory Caching Hypothesis predicts more caches of food f1 in compartment K2 than in compartments K1 or K3.

### 3.5 Analysis

We scored the number of items of each food type that were cached in the three test compartments. Data were collected at the end of each test trial by manually checking the caching trays. Because the specific type of food and location experienced on the first experience trial were counterbalanced among the birds, the predictions about what and where to cache according to the two hypotheses were not identical for all individuals. To overcome this issue, and thus to facilitate the comparison between observed and expected caching patterns, raw data were sorted semantically (Table 1). The three locations were relabelled such that: i) K1 corresponded to the compartment available on experience days 1, 4, 7, and to the hypothetical test on day 10; ii) K2 corresponded to the compartment available on experience days 2, 5, 8, and to the hypothetical trial on day 11; iii) K3 corresponded to the compartment available on experience days 3, 6, 9, and to the hypothetical trial on day 12. Similarly, the two types of food were relabelled such that: i) f1 corresponded to the food received on experience days 1, 3, 5, 7, 9 and on the hypothetical trial on day 11, and ii) f2 corresponded to the food received on experience days 2, 4, 6, 8 and on the hypothetical trial on days 10 and 12.

**Table 1:** Number of items cached in Experiment 1. Raw data and semantically sorted data (K1 = compartment available on trials 1, 4, 7; K2 = compartment available on trials 2, 5, 8; K3 = compartment available on trials 3, 6, 9; f1 = food received on trials 1, 3, 5, 7, 9; f2 = food received on trials 2, 4, 6, 8). The entries in the Sequence column indicate the temporal order of compartments and food type during the familiarisation, e.g. BCA-MP means that the bird was in compartment B on trials 1, 4 and 7 and received macadamia nuts (M) on trials 1, 3, 5, 7 and 9.

| Bird | Sequence | Raw Data |  |  |  |  |  | Sorted Data |  |  |  |  |  |
| --- | --- | --- | --- | --- | --- | --- | --- | --- | --- | --- | --- | --- | --- |
|  |  | A |  | B |  | C |  | K1 |  | K2 |  | K3 |  |
|  |  | P | M | P | M | P | M | f1 | f2 | f1 | f2 | f1 | f2 |
| Washington | BCA-MP | 4 | 2 | 2 | 2 | 3 | 3 | 2 | 2 | 3 | 3 | 2 | 4 |
| Wellington | ABC-MP | 0 | 0 | 0 | 0 | 1 | 0 | 0 | 0 | 0 | 0 | 0 | 1 |
| Caracas | ABC-PM | 8 | 0 | 11 | 0 | 9 | 0 | 8 | 0 | 11 | 0 | 9 | 0 |
| Lisbon | CAB-MP | 1 | 0 | 0 | 1 | 0 | 5 | 5 | 0 | 0 | 1 | 1 | 0 |
| Jerusalem | BCA-PM | 2 | 0 | 3 | 0 | 4 | 1 | 3 | 0 | 4 | 1 | 2 | 0 |
| Quito | CAB-PM | 0 | 0 | 0 | 0 | 0 | 0 | 0 | 0 | 0 | 0 | 0 | 0 |

To determine the factors that best explain the caching behaviour, the number of items *n*_*bfc*_ of food type *f* cached by bird *b* in compartment *c* were treated as random variables drawn for each bird *b* from multinomial distributions with 6 categories (2 × 3 for each food–compartment combination). We compared 13 different possibilities, i.e., models, to set the rates *r*_*bfc*_ of the multinomial distributions. Models with rates independent of food type were characterised by equal rates 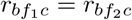 for both food types, whereas in models with food type dependent rates, both 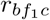 and 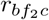 were treated as independent parameters. Similarly, models with rates independent of bird identity had equal rates for all birds 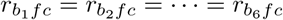 and models with rates independent of compartment had equal rates for all three compartments 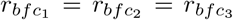. Each model was characterised by the triple (food type dependence, bird identity dependence, compartment dependence), yielding 8 = 2^3^ different models. Effectively, there are only 7 different models, because bird dependence does not matter for compartment and food type independent models, where the rates are equal to 1/6 for each food type and compartment. We used uniform conjugate priors for the Bayesian model comparison. The Future Planning Hypotheses and the Compensatory Caching Hypothesis were formalised with constraints on the prior rates. For prediction 1 of the Future Planning Hypothesis we used a prior with 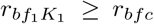 for all *f* ≠ *f*_1_ and *c* ≠ *K*_1_. For prediction 2 of the Future Planning Hypothesis the prior satisfies 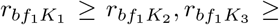 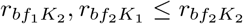 and 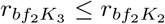. The prior for the Compensatory Caching Hypothesis satisfied the opposite inequalities of prediction 2 of the Future Planning Hypothesis, i.e. 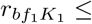 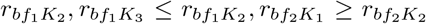 and 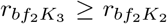. Because each of these three models can be tested with a bird-dependent and a bird-independent variant, we had another 6 models for the Bayesian model comparison such that – together with the 7 models mentioned previously – we tested 13 models in total.

More explicitly, we assume the probability of observing the actual counts *n_bfc_* are given by

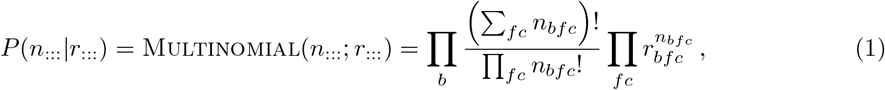

where colons indicate index ranges, e.g. 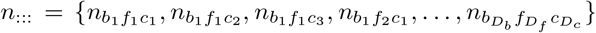, where *D_b_*, *D_f_* and *D_c_* are the numbers of birds, food types and compartments, respectively. If the rates are independent of the food type, i.e. 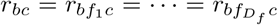, we can rewrite Equation 1 to obtain

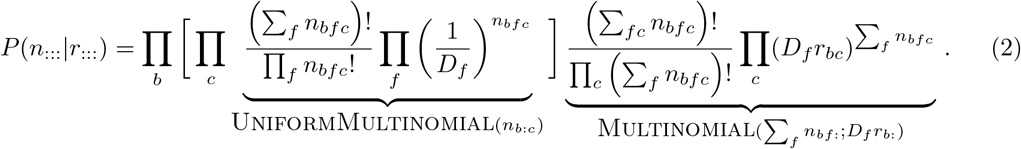

Note that Σ_*c*_ *D*_*f*_*r*_*bc*_ = Σ_*fc*_ *r*_*bfc*_ = 1. Equation 2 has the natural interpretation of a hierarchical data generation process: first, the number of cached items per bird and compartment Σ_*f*_ *n*_*bfc*_ sampled according to rates *D*_*f*_*r*_*bc*_; second, for each bird and compartment the different food types are sampled from a uniform multinomial distribution (in the case of only two food types, from a binomial distribution).

A similar equation can be obtained by rewriting Equation 1 for models with compartment-independent rates. In the case, where the rates are independent of bird identity, i.e. 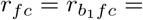 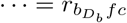, Equation 1 can be rewritten to obtain:

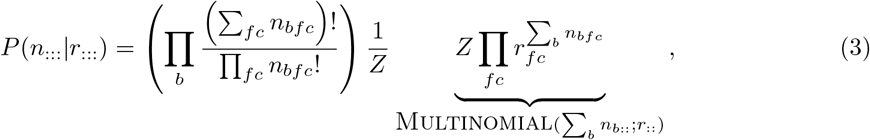

where 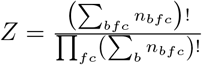.

For the Bayesian model comparison, the posterior *P*(*n*_:::_*|*model) = ∫ *dr*_:::_*P*(*n*_:::_|*r*_:::_)*P*(*r*_:::_*|*model) is obtained by integrating out the parameters weighted by a flat Dirichlet prior *P*(*r*_:::_|model) with parameter *α* = 1. Code and data is available at https://github.com/jbrea/PlanningVsCompensatoryCaching.jl.

### 3.6 Results

Rome and Lima did not eat both types of food in the pre-test in the first pair of trials nor when they were re-tested the second time. Thus, Rome did not proceed to the familiarisation. Although Lima also should not have received the familiarisation, due to an experimenter’s mistake, this bird did receive the familiarisation trials. Neither of these two birds proceeded to the test. In addition, Dublin showed very agitated behaviours during the pre-test, thereby the testing of this bird was stopped on welfare grounds. Consequently, six birds in total passed the criteria of the pre-test and proceeded to the familiarisation. All of these six birds completed the familiarisation and proceeded to the test.

The Bayesian model comparison returns posterior probabilities of the 13 tested models. These posterior probabilities across all models tested sum to 1, and the ratio between the posterior probabilities of any two models would be a Bayes Factor. The comparison revealed that the model in which caching rates depended on bird identity and type of food, but not on compartment, had the highest posterior probability (0.997; Figure 4). In comparison, the highest probabilities for the Future Planning Hypothesis and the Compensatory Caching Hypothesis were 0.0014 and 0.00001, respectively (red dashed bars for FPH 1 and CCH in Figure 4). The difference in the posterior probability of the Future Planning Hypothesis and the Compensatory Caching Hypothesis results from the caching pattern of a single bird: Lisbon cached as expected under the Future Planning Hypothesis. If Lisbon’s data is excluded from the model comparison, the posterior probability of the Compensatory Caching Hypothesis is higher (0.0002) and the posterior of the Future Planning Hypothesis is lower (0.00008) In any case, neither the Future Planning nor the Compensatory Caching hypotheses explained the obtained data well. Instead, the data seemed to be best explained by individual preferences for different foods by the different birds.

**Figure 4:**
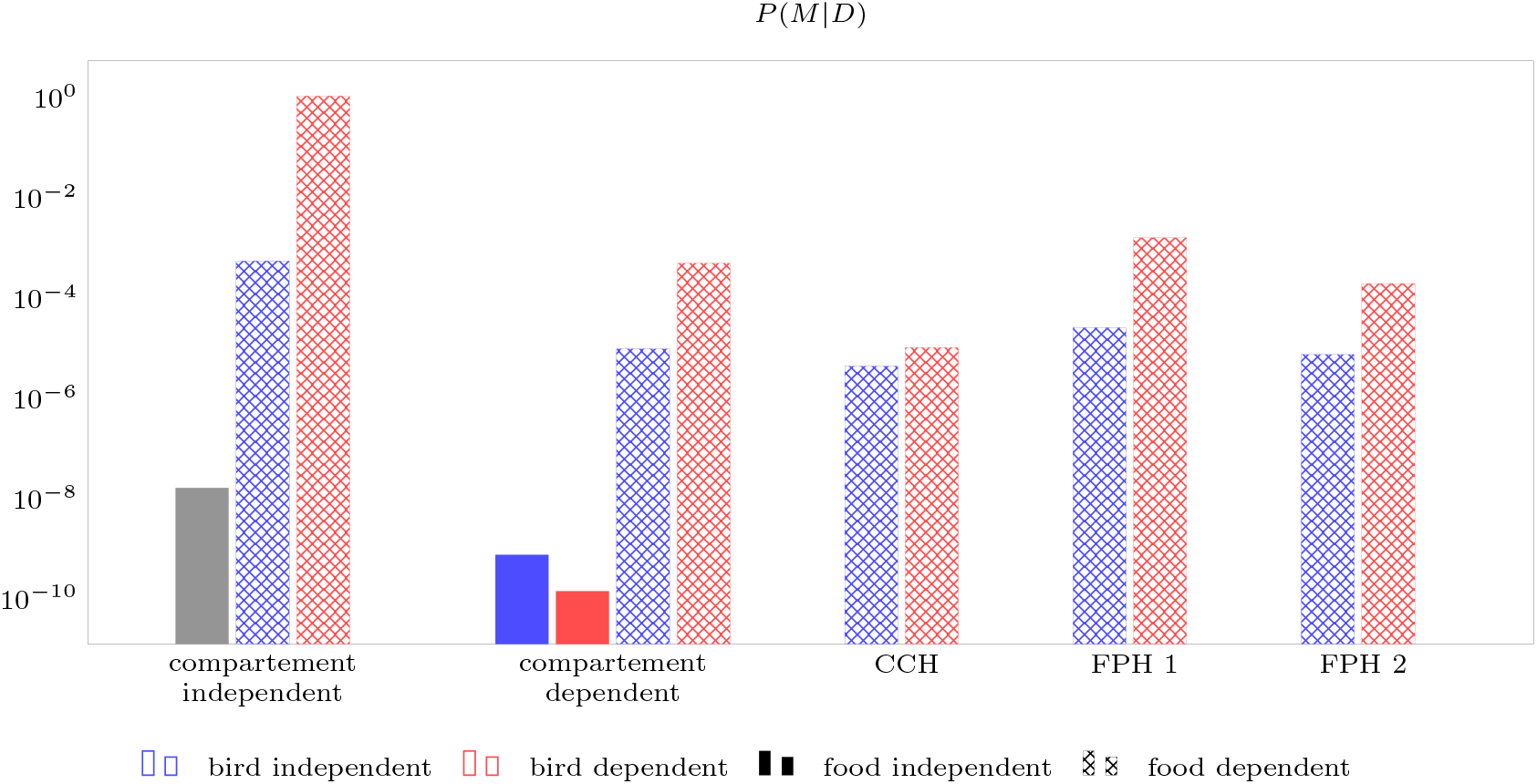
**The data is best explained by a model where the caching rate depends on the bird identity and the food type but not on the compartment** (red dashed bar at compartment independent). The Future Planning Hypotheses (FPH 1 and FPH 2) and the Compensatory Caching Hypothesis (CCH) have much lower posterior probability because they rely on compartment- and food-dependent caching rates. Because the compartment- and food-type-independent models assume that the caching rate for each of the three compartments and food types is 1/6, there is no difference between a bird-dependent and a bird-independent model (gray bar).

In summary, the overall pattern of results did not provide support for either the Compensatory Caching Hypothesis or the Future Planning Hypothesis. One possible explanation for this inconclusive outcome may be that the cognitive load imposed on the jays in this test was too high, thereby impairing their ability to maximise caching outcomes according to future scenarios or food-specific associations with the different locations. To test between the two hypotheses in a scenario with a decreased cognitive load, we conducted a second experiment. Here, the cognitive load was decreased through a different manipulation of food availability: instead of having access to one of two foods on alternating days during the experience phase, birds now either had access or no access to one type of food.

## 4 Experiment 2

Experiment 2 employed the same protocol as that used in Experiment 1, but with one key difference: to decrease the cognitive load associated with the task, only one type of food was used. The type of food was not the same for all birds, but was either peanuts or macadamia nuts, based on what food the bird preferred. Like in Experiment 1, the test compartment that was accessible rotated on a 3-day basis. Birds either received food (Food condition, F) or received no food (No Food condition, N) on alternating days during the experience phase. This food availability rotated on a 2-day basis, i.e. food was available on every other trial. This experiment was pre-registered on the Open Science Framework (https://osf.io/3y5tm/). The pre-registration was finalised after the pre-tests were completed but before the start of the test, i.e., prior to the first experience trial. The experiment was conducted in October 2018. PA was the experimenter.

### 4.1 Pre-Test 1

Pre-test 1 was conducted to ascertain individual preferences for caching two different types of food. This information was used to decide which food would be provided to each individual bird in the test to minimise the probability that a bird would not cache during the test and thus minimise a decrease of the sample size. Approximately 1.5h after their maintenance diet had been removed from the aviary, a bird was tested in visual isolation from the rest of the group, namely in an indoor compartment that was not part of the experimental set-up. The bird was provided with one caching tray and with two food bowls, one containing 50 peanuts and the other containing 50 macadamia nut halves. The bird was allowed to eat and manipulate (including caching in the compartment) the food for 15 minutes. At the end of the trial, the bird was released back into the outdoor aviary. The experimenter manually checked the caching tray and recorded the number of items of each type of food cached. Approximately 3h after each trial, the bird was again given access to the same compartment and given the opportunity to retrieve its caches for 10 minutes. Retrieval sessions were conducted only to minimise the probability that birds would stop caching because this behaviour yielded no benefit to them. Each bird received three trials in total. Individual preferences were established by considering – for each bird separately – the type of food that was cached on average more often across the trials. To proceed to pre-test 2, a bird was required to have cached at least one food item in at least two of the three trials.

### 4.2 Pre-test 2

Eurasian jays usually need repeated exposure to a food to start eating it in a novel situation, even if they had previously received the same food. Given that the jays had not had the opportunity to consume powdered food following the completion of Experiment 1 (i.e. from January to October 2018), it was necessary to ensure that the birds were still motivated to eat powdered food as this food was subsequently used in the test. To this end, pre-test 2 was conducted to ascertain that birds would eat peanuts and macadamia nuts in powdered form.

Pre-test 2 involved two stages. In stage 1, birds were given the opportunity to eat both types of powdered food in the aviary as a group. Powdered peanuts and macadamia nuts were presented in separate bowls, after the birds’ maintenance diet had been removed from the aviary for approximately 1.5 h. Birds were observed by the experimenter from an observation hut adjacent to the aviary. If higher-ranking individuals monopolised the food bowls, they were separated from the group, such that lower ranking individuals could access the food. The experimenter scored the number of times each bird inserted its beak into each food bowl. To proceed to stage 2, a bird was required to insert its beak at least twice into their preferred food (as determined in pre-test 1). If this criterion was not reached by all birds on the first trial, the trial was repeated up to two times (i.e. a maximum of three trials in total). In stage 2, a bird was tested individually in an indoor compartment that was not part of the experimenter set-up. Like in stage 1, birds were tested after their maintenance diet had been removed from the aviary for approximately 1.5h. The bird was allowed to eat freely from a bowl containing 50.0 g of its preferred food in powdered form for 15 minutes. A bird was required to eat at least 0.1 g of food to proceed to the familiarisation. If a bird did not pass this criterion, the trial was repeated up to two times (i.e. a bird received a maximum of three trials in total).

### 4.3 Familiarisation

The Familiarisation followed the same procedure as described in Experiment 1.

### 4.4 Test

The test followed the same procedure as in Experiment 1. However, instead of two foods, birds received only one food, i.e. their preferred food. This procedure was chosen to maximise the probability that each bird would eat the food during the test. Again, the experiment consisted of an experience phase involving nine experience trials, followed by one test trial. In the experience phase birds received one trial per day on nine consecutive days. Each trial was conducted in the morning, after the birds’ maintenance diet had been removed from the aviary for approximately 1.5 h. On each trial a bird was individually allowed to access the experimental set-up through the trap-window in the middle compartment and was given access only to one of the three test compartments. On the platform was either a bowl containing 50.0 g of its preferred food in powdered form (Food condition, F) or an empty bowl (No Food condition, N). In the Food condition, food was available in powdered form to prevent the bird from caching in the test compartments during the experience phase. In the No Food condition, an empty bowl – rather than no bowl at all – was placed in the accessible compartment, both to ensure that it was only the availability of food that differed between the two conditions and to maximise the probability that the bird would enter the testing compartment when no food was available. The bird could freely move between the middle compartment and the accessible test compartment and – in the Food condition – eat the food for 15 minutes. At the end of a trial, the bird was released into the outside aviary through the trap-window in the middle compartment. Birds were tested in the same order on all trials. Like in Experiment 1, the location and availability of food experienced by the birds on each trial changed according to two different patterns (Figure 5). The test compartment that was accessible rotated on a 3-day basis, so that the birds that had access to compartment A on trial 1 experienced the test compartments in the following order across the nine trials: A-B-C-A-B-C-A-B-C. The availability of food rotated on a 2-day basis, i.e. food was available every other trial. As a result, all test compartments were experienced by each bird three times, and differed in the frequency in which they were associated with the food. Specifically, two compartments were associated twice with the presence of food and once with the absence of food, whereas the third compartment was characterised by the opposite pattern. The compartment that was accessible on trial 1 and food availability were counterbalanced across birds. This meant that – as far as food availability is concerned – there were two groups of birds: one group that experienced food on trial 1 (henceforth Food-First group, FF) and the other group that experienced no food on trial 1 (henceforth Empty-First group, EF).

**Figure 5:**
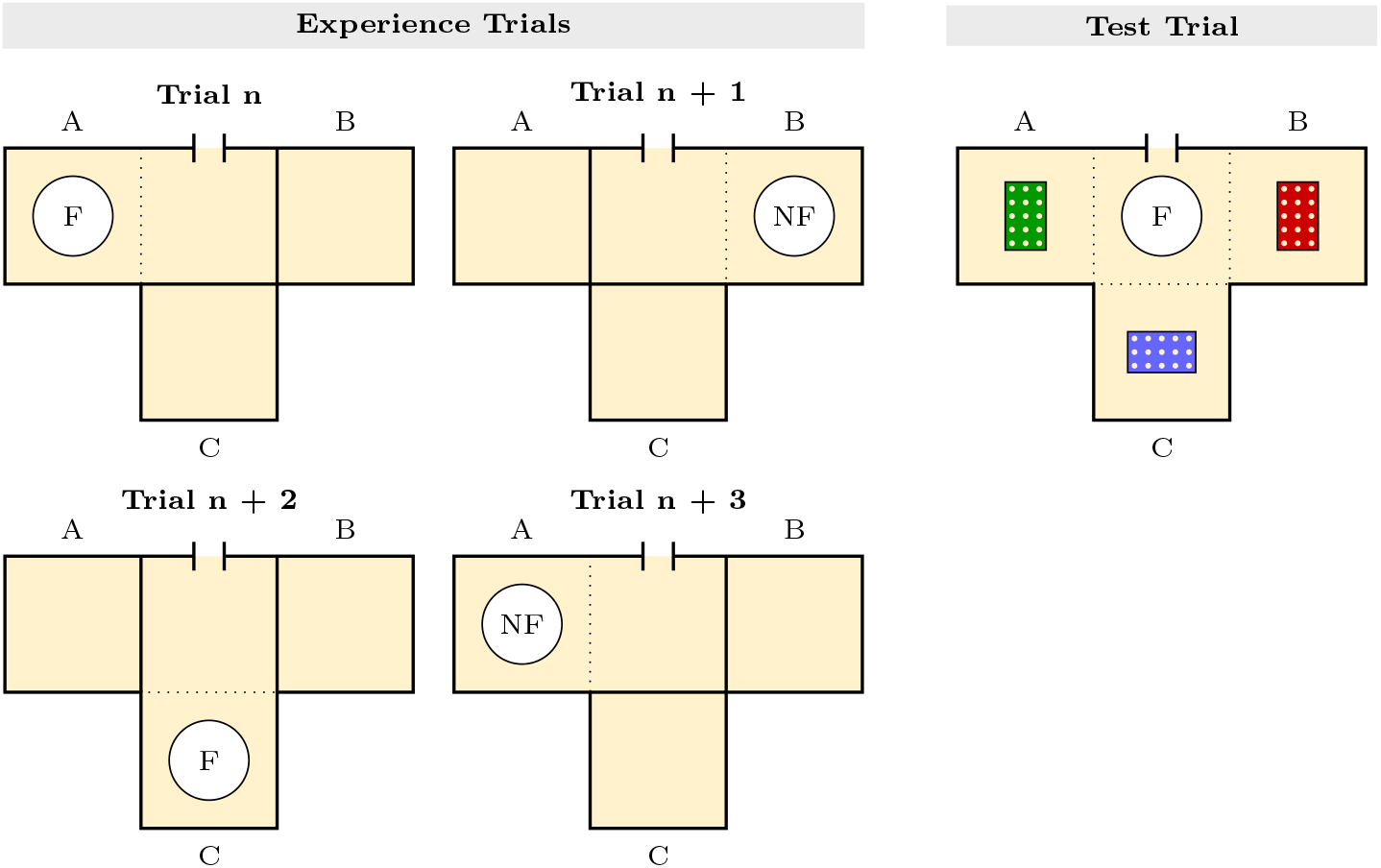
Top view of the experimental set-up of Experiment 2. Birds could access the three test compartments (A, B, C) from a middle compartment. On each experience trial (Left), the birds had access only to one test compartment (e.g. A in Trial n; B in Trial n+1), and they were presented either with a bowl containing powdered food (F, Trial n, Trial n+2) or with an empty bowl containing no food (NF, Trial n+1, Trial n+3). The compartment that was accessible and food availability rotated on a 3-days cycle and on a 2-days cycle, respectively. On the test trial (Right), a bowl containing food items (i.e. not food in powdered form) was placed in the middle compartment and caching trays were placed in all test compartment. Birds could freely move and cache in all compartments.

The test phase was conducted on the day of the last experience trial (9th day), approximately 3h after the experience phase. Again, a bird was individually given access to the experimental set-up. All test compartments were accessible at this stage and each contained one caching tray (5 × 3 seedling pots filled with sand). A bowl containing 50 items of the bird’s preferred food (whole peanuts or macadamia nut halves) was placed in the middle compartment. The bird could eat and cache the food in all test compartments for 15 minutes. At the end of the trial, the bird was released into the outdoor aviary through the middle compartment. Birds were not given the opportunity to retrieve their caches.

### 4.5 Predictions

According to the Compensatory Caching Hypothesis birds should cache a higher proportion of food items in the compartment(s) where – during the experience phase – the food had been available only once than in the compartment(s) where the food had been available twice. Importantly, birds in the Food-First group experienced two compartments as being more frequently associated with food, whereas birds in the Empty-First group experienced only one compartment as being more frequently associated with food. As a result, the Compensatory Caching Hypothesis predicts that birds in Food-First group should cache more in one compartment, but that birds in the Empty-First group should concentrate their caches equally across two compartments (Figure 6).

**Figure 6:**
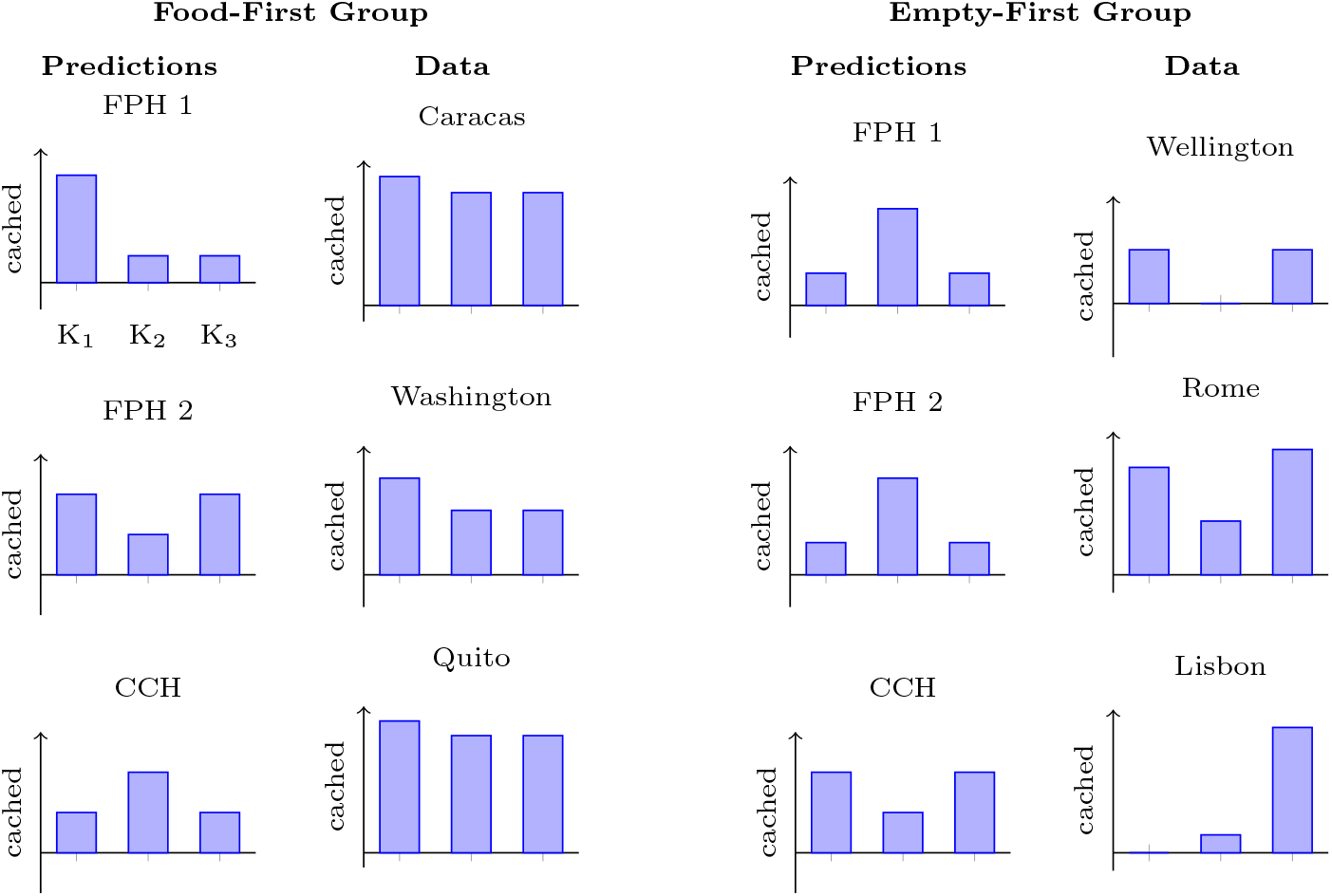
Individual caching patterns allow no conclusion. A bird in the Food-First group should expect not to receive powdered food on test day 10. If it caches according to the prediction 1 of the Future Planning Hypothesis (FPH 1), it will provision only for the next day, where it expects to be in compartment K1 with no food available; thus it would cache most in K1. If it takes into account the next three days (FPH 2), it will distribute the caches in compartment K1 and K2. If the bird caches according to the Compensatory Caching Hypothesis (CCH), the bird should cache more in compartment K2, where it has only once encountered the powdered food. The predicted caching pattern for birds in the Empty-First group is reversed, because on day 10 it will have food available in compartment K1. The two variants of the Future Planning Hypothesis are indistinguishable for the Empty-First group. The actual numbers in the predictions serve illustration purposes only. The hypotheses allow only qualitative predictions, e.g. the Compensatory Caching Hypothesis predicts more caches in compartment K2 than in compartments K1 or K3 for birds in the Food-First group.

By contrast, according to the Future Planning Hypothesis birds should cache more in the compartment(s) in which they expect that they would receive no food, either on the next day (hypothetical experience trial 10, prediction 1) or on the following three days (hypothetical experience trials 10 to 12, prediction 2). Both of these predictions are applicable to the Food-First group (Figure 6). However, only prediction 2 is relevant to the Empty-First group. During the nine experience trials, these birds were given food every other day in the following sequence: N-F-N-F-N-F-N-F-N, such that according to this pattern they would receive food on the hypothetical trials 10 and 12 but not on trial 11. As a result, if jays can cache according to future need, then it makes little sense for birds in the Empty-First group to cache in the compartments where they expect to be on day 10 (prediction 1) because there, food would be available anyway. Rather they should concentrate their caches to the compartment to which they expect to be given access on day 11 because it is this compartment only in which no food would be available across the three hypothetical trials (prediction 2).

### 4.6 Analysis

Data were collected, scored and semantically sorted by using K1-K3 as variables as in Experiment 1 (Table 2). The analysis was the same as in Experiment 1, except for the following differences. To determine the factors that best explain the caching behaviour, the number of items *n_bc_* cached by bird *b* in compartment *c* were treated as random variables drawn for each bird *b* from multinomial distributions with 3 categories (one for each compartment). We compared 9 different possibilities (models) to set the rates *r_bc_* of the multinomial distributions. Each model was characterised by the tuple (bird identity dependence, compartment dependence), yielding 3 = 4 − 1 different models; there is only one compartment-independent model with rates 1/3 for each compartment. Six additional models with constrained prior rates were tested to quantify the posterior probabilities of the Future Planning Hypothesis (predictions 1 and 2) and the Compensatory Caching Hypothesis, for both bird-dependent and bird-independent rates. The constraints on the prior rates were 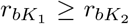 and 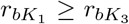 for prediction 1 of the Future Planning Hypothesis for birds in the Food-First group, 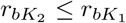 and 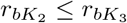 for prediction 2 of the Future Planning Hypothesis for birds in the Food-First group and for the Compensatory Caching Hypothesis for birds in the Empty-First group, 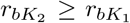 and 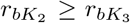 for the Compensatory Caching Hypothesis for birds in the Food-First group and for the Future Planning Hypotheses (both variants) for birds in the Empty-First group.

**Table 2:** Number of items cached in Experiment 2. Raw data and semantically sorted data (K1 = compartment available on trials 1, 4, 7; K2 = compartment available on trials 2, 5, 8; K3 = compartment available on trials 3, 6, 9). The entries in the Sequence column indicate the temporal order of compartments and conditions during the familiarisation, e.g. BCA-NF means that the bird was in compartment B on trials 1, 4 and 7 and received no powdered food (N) on odd days and powdered food (F) on even days.

| Bird | Sequence | Raw Data |  |  | Sorted Data |  |  |
| --- | --- | --- | --- | --- | --- | --- | --- |
|  |  | A | B | C | K<sub>1</sub> | K<sub>2</sub> | K<sub>3</sub> |
| Washington | CAB-FN | 2 | 2 | 3 | 3 | 2 | 2 |
| Wellington | CAB-NF | 0 | 1 | 1 | 1 | 0 | 1 |
| Caracas | BCA-FN | 7 | 8 | 7 | 8 | 7 | 7 |
| Rome | ABC-NF | 6 | 3 | 7 | 6 | 3 | 7 |
| Lisbon | BCA-NF | 7 | 0 | 1 | 0 | 1 | 7 |
| Quito | ABC-FN | 9 | 8 | 8 | 9 | 8 | 8 |

### 4.7 Results

All birds passed the pre-tests and the familiarisation, except for Dublin who did not successfully complete the familiarisation. Therefore, seven birds in total took part in the test. On the ninth and last experience trial, Lima went through a small empty space in between the suspended platform in the compartment and the mesh wall and therefore experienced not only the accessible compartment but also another one. Thus, testing of this bird was stopped, i.e. Lima did not receive the test trial. Thus, only six birds were tested in the test trial. All birds except one (Rome) had participated in the test trial of Experiment 1.

As in Experiment 1, the posterior probability of the model with compartment independent caching rates was highest (0.72; gray bar in Figure 7). The posterior probability of the Compensatory Caching Hypothesis (0.16; blue bar at CCH in Figure 7) is higher than those of the Future Planning Hypothesis (0.002; blue bars at FPH 1 and FPH 2 in Figure 7). This is consistent with the observation that the caching patterns of Wellington, Rome and – to some extent – Lisbon are compatible with the Compensatory Caching Hypothesis, whereas Caracas, Washington and Quito distributed their caches almost equally among the three compartments.

**Figure 7:**
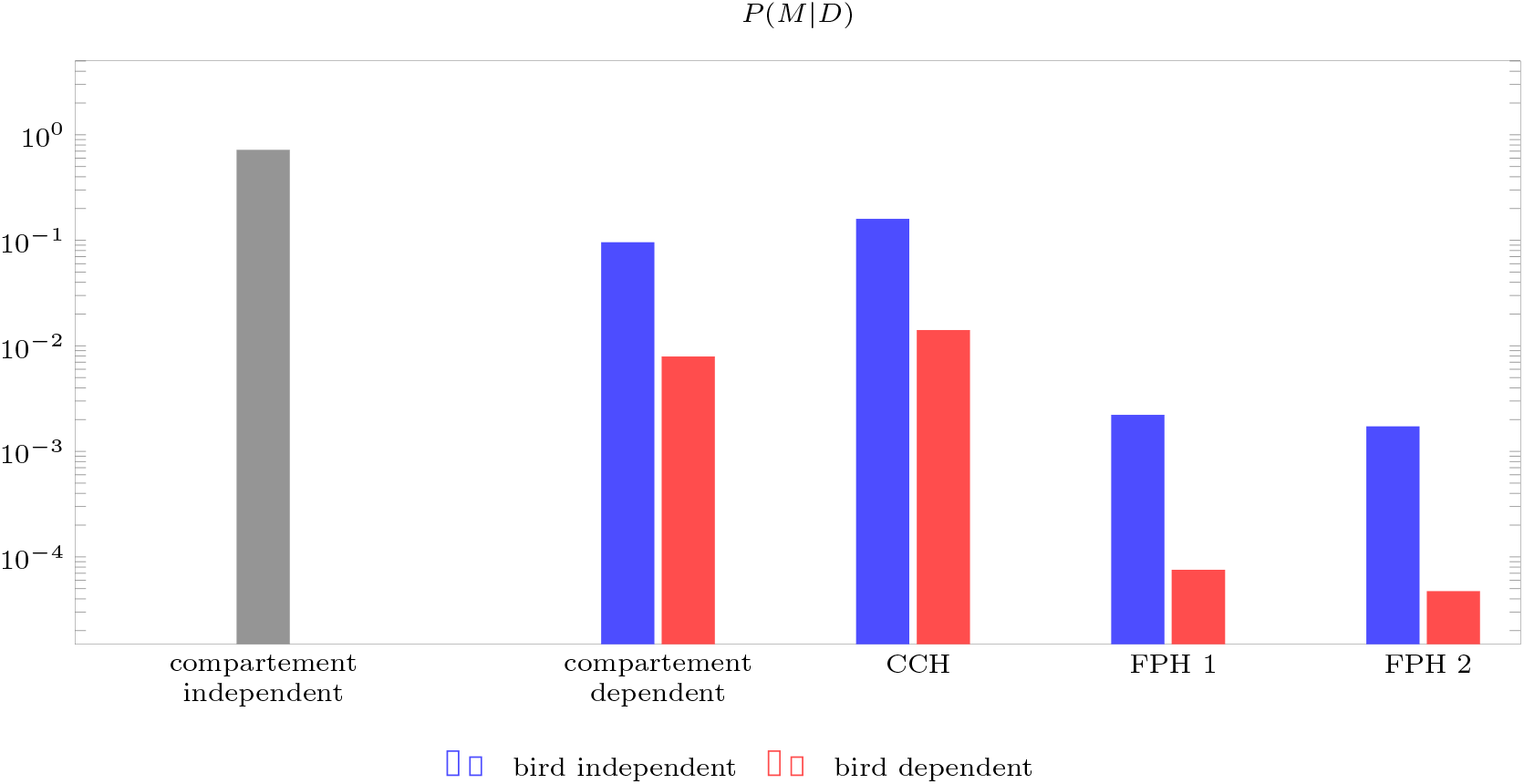
The data is best explained by a model where the caching rate does not depend on the compartment (gray bar at compartment independent). The Future Planning Hyptheses (FPH 1 and FPH 2) and the Compensatory Caching Hypothesis (CCH) have much lower posterior probability because they rely on compartment-dependent caching rates. Because the compartment-independent model assumes that the caching rate for each of the three compartments is 1/3, there is no difference between a bird-dependent and a bird-independent model.

In summary, even though the Compensatory Caching Hypothesis had a higher posterior probability than any variant of the Future Planning Hypothesis, we found that, as in Experiment 1, an equal distribution of caches among the three compartments explained the data best.

## 5 Discussion

The aim of this study was to test two different hypotheses regarding the suggested ability of corvids to maximise caching outcomes according to future scenarios, namely the Compensatory Caching Hypothesis and the Future Planning Hypothesis. To this end, we tested Eurasian jays in two experiments, for which the two hypotheses have opposite predictions regarding the jays’ behaviour. Across both experiments, the data did not support either hypothesis: the model with the best fit to the data was dependent on the identity of the bird and, in Experiment 1 (where there was a choice of food types), also the type of food cached. The birds cached predominantly their preferred food across the different locations regardless of their previous experiences in those locations.

Thus, the current results may seem to contrast the interpretation of results of previous studies, in particular the results reported by Raby et al. [9], the study that our experimental design was based on. Raby et al. [9] reported that California scrub-jays preferentially cached food where it would not be available in the future; thus, these results would appear to show a caching pattern that is – unlike the data from the current study – dependent on the location and the experimental manipulations experienced by the birds in the different locations. Given that the experiments reported here and those conducted by Raby et al. [9] employed different methods, it is not possible to draw conclusive inferences about what factors may have caused inconsistent results between the two studies. A crucial step for future research is to replicate the findings reported here in Eurasian jays, and those obtained by Raby et al. [9] in California scrub-jays. Subsequently, in case the results of both studies will be corroborated, future work should investigate the factors (e.g. experimental paradigms, model species) that may have caused inconsistent outcomes between these two studies.

One factor that future research may focus on is whether methodological differences in housing condition may have played a role. Specifically, the scrub-jays tested by Raby et al. [9] were housed in pairs in 2 × 1 × 1 m cages. However, during testing (experience and test trials) only the focal individual had access to the testing compartments. On the other hand, the Eurasian jays that participated in the current study were housed as a group, and they were tested serially in the same indoor compartments, i.e., each bird had access to the same testing compartments on each day. This means that, in contrast to the scrub-jays in Raby et al. [9], the Eurasian jays tested here could see other individuals having access to the compartment that they were also being tested in throughout the experience and test trials.

Another potentially relevant factor to consider is whether differences in the experimental paradigms may have influenced the ecological validity of the tasks, and consequently affected jays’ capability to cache according to future needs. A key difference between the studies is that the association between food and locations was stable across experience trials in Raby et al. [9]’s experiments but it changed systematically in the experiments reported here. It is possible that the ‘dynamic’ food-location association in Experiments 1 and 2 may have impaired the ecological relevance of the task. Eurasian jays are specialised cachers that primarily rely on trees’ nuts (i.e. acorns) as food for caching [27, 30]. Thus, it may be unlikely that a wild Eurasian jay would experience locations in which a specific food is abundantly available or absent in a cyclic fashion with changes occurring over very short time intervals: a nut tree does not bear abundant fruits every few days but no fruits in between. Therefore, future work on the cognitive mechanisms underpinning caching behaviour in corvids may want to evaluate the interplay between experimental designs and ecological validity of the task. To this end, it may also be beneficial for both theory development and experimental design to take into account species-specific differences in caching given that these features may influence the ecological relevance of the tasks for different corvids.

The difference in experimental design between the Raby et al. [9]’s study and the current study noted above further highlights the fact that the current study likely has an increased cognitive load due to the rapid changes of food availability between three different locations. However, in their current form, neither the Compensatory Caching Hypothesis nor the Future Planning Hypothesis involve any auxiliary claims regarding the extent of information that the underlying mechanism is able to process. The Future Planning Hypothesis specifies that individuals should be able to keep track of ‘what-where-when’ representations to select actions that are most appropriate at a given point; thus, it appears that the cognitive load involved in this study, namely to integrate information about the food availability at different locations based on time intervals, is compatible with the hypothesis. Similarly, the Compensatory Caching Hypothesis specifies that the suitability of a given caching location for a specific food is based on food-specific ‘weights’ associated with each location, which are updated when the bird finds itself at that location. Thus, in its current form the Compensatory Caching Hypothesis also seems compatible with the cognitive load involved in the current experiments. Future theoretical work may, however, expand the hypothesis regarding the manner in which the ‘weights’ are updated, and thus should motivate further empirical work.

A final consideration should be made with regard to sensitivity of our study. While our results do not provide strong support for either the Future Planning Hypothesis or the Compensatory Caching Hypothesis, it is also possible that our experimental design may have lacked the sensitivity to detect some effect sizes that are compatible with both hypotheses. Specifically, as we had only a single test trial for each of the 6 birds that completed both experiments, it is plausible that the general variability of our Eurasian jays’ caching behaviour is too large for effects consistent with either hypothesis to be regularly detected in our design. Nevertheless, our results are inconsistent with the possibility of very large future planning or compensatory caching effects in our design.

In summary, the data from this study provide no support for either of the two tested cognitive abilities being the base of future-oriented caching behaviour in Eurasian jays. To evaluate whether, by contrast, a third explanation that has been proposed in the literature and discussed in the introduction, namely the Mnemonic Association Hypothesis [17], could provide a better explanation for the Eurasian jays’ caching behaviour, future studies could utilise equivalent procedures that can simultaneously test between different competing hypotheses.To increase the sensitivity of the study, future experiments may wish to develop designs in which large raw effect sizes are predicted by each hypothesis, or designs in which multiple testing trials can be performed by each bird. In addition, future studies repeating the experiments reported here both in Eurasian jays and other corvid species are needed to strengthen the claims that can be made based on the current findings. Ultimately, this line of research will likely play a key role in elucidating the cognitive underpinnings of caching decisions in corvids.

## Ethics

All procedures were approved by the University of Cambridge Animal Ethics Review Committee.

## Data accessibility

Data and code are available at https://github.com/jbrea/PlanningVsCompensatoryCaching.jl.

## Authors’ contribution

JB and LO designed the study. PA, LO, and BGF collected the data for Experiment 1 and PA collected data for Experiment 2. JB analysed the data. PA wrote the first draft of the manuscript with critical inputs from JB and LO. NSC and BGF provided helpful comments on subsequent drafts, and all authors commented on the final draft.

## Acknowledgements

JB thanks Alireza Modirshanechi for helpful discussions and feedback on the Bayesian model comparison.

